# Effect of a LINE1 DNA sequence on expression of long human genes

**DOI:** 10.1101/2023.11.21.568109

**Authors:** Jay C. Brown

**Affiliations:** Department of Microbiology, Immunology and Cancer Biology University of Virginia School of Medicine, Charlottesville, Virginia, United States of America

## Abstract

The study described here was carried out to pursue the idea that a truncated, transposition incompetent fragment of a LINE1 retrotransposon may affect the expression of a human gene when it is located inside the gene sequence. NCBI BLAST was used to probe the human genome to identify protein coding genes containing an abundant ∼1500bp LINE1 fragment (called t1519) in the gene body. The length and expression level of such genes was then compared with the same properties in genes that lack t1519 in human chromosomes 16-18. The results showed a striking effect of t1519 on long genes, those with lengths greater than ∼140 kb. Nearly all were found to have one or more t1519 sequences in the coding region. In contrast, genes in the common length range (less than 140 kb) could either have t1519 or not. A correlation was also observed with the level of gene expression. While expression of long, t1519-containing genes was limited to ∼50 TPM, genes in the common length range could be much higher, in the range of 500-600 TPM, regardless of whether or not they have t1519 elements. Contrasting results were obtained when the analysis was performed with lncRNAs rather than with protein-coding genes. Among lncRNA genes a chromosome-specific effect was observed. Restricted expression correlating with the presence of t1519 was observed in both long and common length genes of chromosomes 16 and 17, but not in chromosome 18. The results are interpreted to support a strong suppressive effect of t1519 on expression of long protein coding genes and on both long and common length lncRNA genes of chromosomes 16 and 17. It is suggested that the suppressive effect on expression, particularly among long genes, meets a need for the cell to limit the overall level of transcription it can support.

**Author summary:** Although LINE1 DNA sequence elements are well known for their ability to replicate and move autonomously within the human genome, these features are observed in only a small proportion (0.02%) of the total human LINE1 population. Nearly all of the total ∼500,000 LINE1 elements are fragments of full-length LINE1 and are inactive for autonomous replication or movement. Truncated, inactive LINE1 sequences are found throughout the human genome including within the body of protein-coding genes, and this intragenic population is the subject of the study described here. The goal was to extend what is known about the properties of intragenic LINE1 sequences. The study was carried out with t1519, a truncated LINE1 sequence composed of the 3’ terminal ∼1500 bp of the ∼6000 bp full length LINE1 element, and with the sequences of three human chromosomes 16, 17 and 18, that are rich in t1519 sequences. NCBI BLAST was used to identify t1519-containing genes in each chromosome, and the length and expression level of those genes was compared with control genes lacking t1519. A striking result was observed in the case of long protein-coding genes, genes longer than 140 kb. Nearly all had one or more t1519 sequences in the gene body, all in introns. An effect on the level of gene expression was also observed. Low expression (<50 TPM) was found in all long, t1519 positive genes while much higher levels (500-600 TPM) were found with genes in the common length range (< 140 kb) regardless of the presence of t1519. Similar results were obtained when lncRNA genes were studied instead of protein-coding ones. The results are interpreted to support a strong suppressive effect of t1519 on expression of long protein coding genes and also on certain lncRNA genes. It is suggested that the suppressive effect is due to a need for the cell to limit the overall level of transcription it can support.

## Introduction

LINE1, non-LTR retrotransposons are well-known for their ability to replicate autonomously and move within the genomes of plant and animal species including humans. Movement and replication-competent human LINE1 (L1) elements are ∼6000 bp in length and it is estimated that there are up to 100 that are currently active for transposition. These are far outnumbered, however, by the ∼500,00 truncated forms of L1 that are not active for transposition but constitute 16.9% of the human genome [1, 2]. It is natural to be curious about the function of this large fraction and clarifying the function is the goal of the investigation described here.

Prior studies of truncated L1 function have suggested these elements have a role in reducing the expression of genes that contain them [3–7]. Suppression is proposed to be due to the fact that both full-length and truncated L1s are rich in polyadenylation sites such as AATAAA and ATTAAA that terminate transcription. Experimental results suggest that by causing premature termination of transcription, L1 polyadenylation sites may act to reduce the expansion of L1s in the genome and also suppress expression of L1-containing genes in the “host” cell.

The strategy employed to clarify truncated L1 function was to begin with a single, abundant L1 element and then use NCBI BLAST to identify human genes that contain the element inside a protein coding gene. The lengths and expression levels of the genes identified were then compared with a population of control genes lacking the L1 sequence with the hope that the comparison would be revealing about L1 function.

Results of the analysis have indicated the involvement of long genes, genes greater than 140 kb in length and accounting for ∼10% of the human genome. Due to their greater length, long genes require a longer time to be transcribed by RNA polymerase and they are exposed to other transcriptional delays due, for instance, to transcriptional pauses and template DNA overwinding [8–11]. Studies of long gene tissue distribution have revealed they are concentrated in genes of the nervous system where they encode proteins involved in cell adhesion, ion channels and receptor function among others [12–13]. Long genes are thought to have arisen evolutionarily at least 320 million years ago as they are found in the DNA sequence of the green anole lizard and other species that have existed since then [13–14]. Studies tracing the evolution of long genes have shown they develop from shorter gene homologs mainly by expansion of the introns [15–16].

## Materials and methods

The t1519 sequence was identified in a pairwise DNA sequence comparison of two human genes, CFHR1 (chromosome 1) and TSHR (chromosome 14). The comparison was performed with Matcher from Job Dispatcher available from the European Bioinformatics Institute (https://www.ebi.ac.uk/jdispatcer) The location of t1519 within LINE1 was identified by aligning the t1519 sequence with human retrotransposon AF148856. Phylogenic analysis of the t1519 nucleotide sequence was performed with Clustal Omega also from the European Bioinformatics Institute (https://www.ebi.ac.uk/jdispatcher/msa/clustalo) The sequence of t1519 can be downloaded from Supplementary Table S1.

NCBI BLAST was used to scan the human genome (taxid 9606) with the t1519 DNA sequence using the option for highly similar sequences. Two features of the output were saved: (1) the number of matches in each chromosome; and (2) the location of each match. Match locations were saved as .bed files and viewed with Integrated Genome Viewer (https://igv.org/app/) Bed files can be downloaded from Supplementary Tables S2 S3 and S4 for chromosomes 16, 17 and 18, respectively. Human genome build hg38 was used for chromosomes 16 and Y; CHM13 was used for chromosomes 17, 18 and 20.

Data for plots of gene expression against length were obtained from the GTEx Portal (https://www.gtexportal.org/home/) and UCSC Genome Browser (https://genome.ucsc.edu), respectively. Data for plots of t1519 count against gene length were obtained from this study and from UCSC genome Browser, respectively. In both cases the results were saved with Excel (see Supplementary Tables S5-10) and plotted with SigmaPlot 15.0.

## Results

### Identification of an appropriate truncated L1 sequence

For use in the experimental strategy described above, it was considered that an ideal truncated L1 probe would be a single L1 region rather than a group of L1 sequences. The probe should also be widely distributed in the human genome so that a large number of probe-containing genes would be identified. Consideration of the available options led to the adoption of t1519, a sequence composed of the rightmost 1519 bp of a canonical, ∼6000 bp L1 sequence (Fig. 1). t1519 contains a substantial portion of the L1 ORF2 gene that encodes a reverse transcriptase involved in L1 transposition. It also encodes five polyadenylation transcription termination sites, two involved in termination of L1 ORF2 transcription and three others with unknown functions [4]. Phylogenetic analysis of the t1519 DNA sequence showed a high degree of homology to L1 sequences of gorilla and chimpanzee. Homology with mouse L1, however, is weak (Fig. 2).

**Fig. 1:**
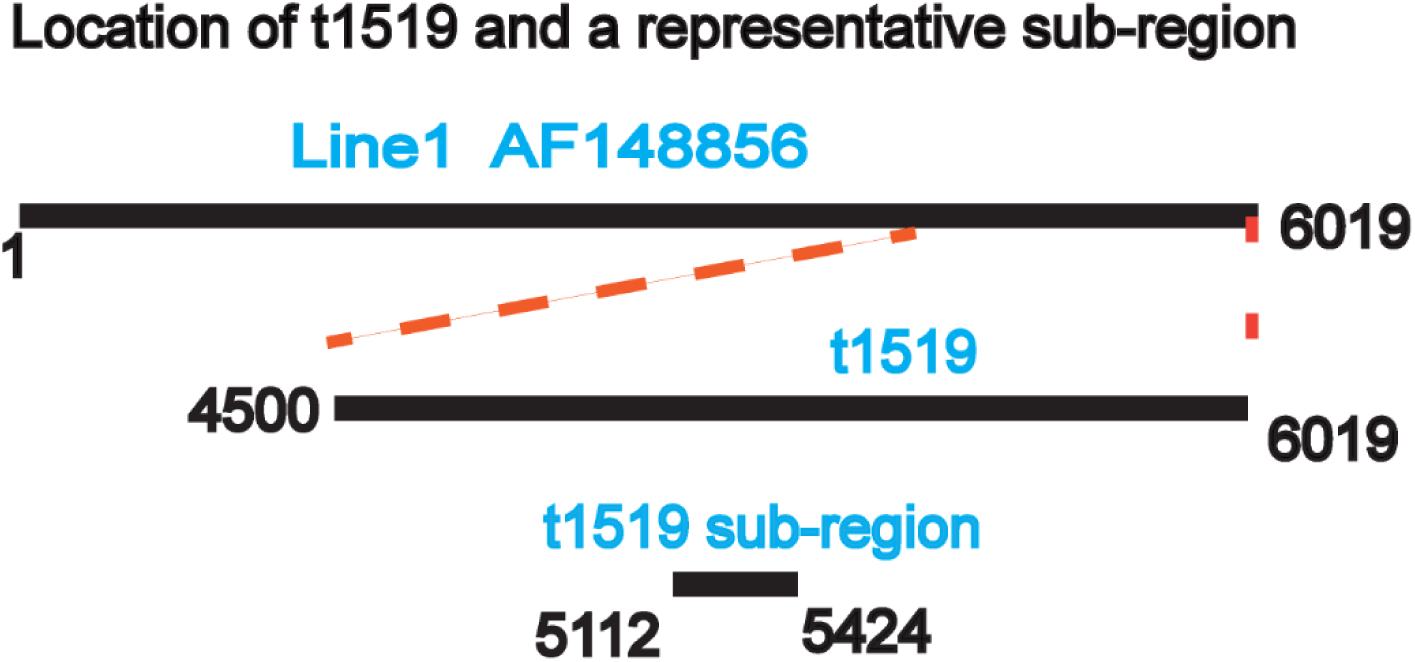
Diagram showing a full-length L1 element, the region of L1 present in t1519, and a sub-region of t1519. The diagram illustrates that some of the sequences returned by NCBI-BLAST scanning with t1519 are not full-length t1519 but sub-regions such as that illustrated here.

**Fig. 2:**
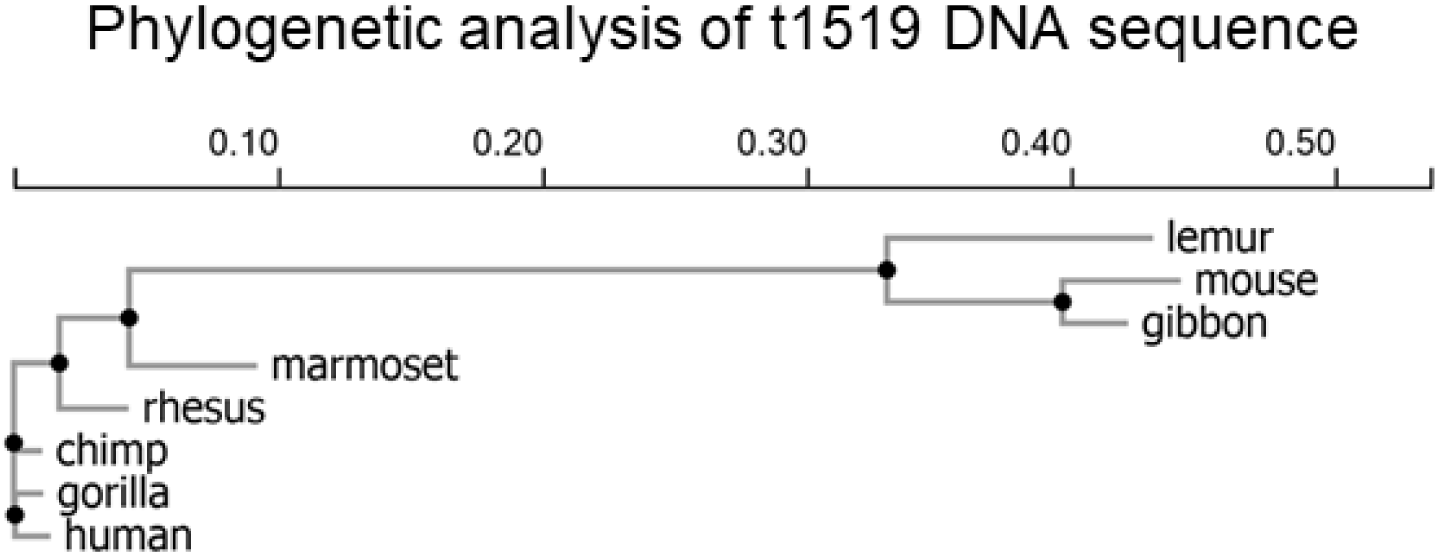
Phylogenetic analysis of the t1519 DNA sequence compared to homologs in other species. Note the close relationship of the human sequence with chimpanzee and gorilla sequences.

### BLAST scan of the human genome with t1519

BLAST analysis was performed with t1519 sequences in two forms: (1) the entire t1519 sequence, and (2) eight sub-regions of the full-length t1519 sequence each 180 bp in length. These were numbered regions 1-8 beginning at the 5’ end of the t1519 sequence. Table 1 shows the number of t1519 matches revealed in the scan. The results demonstrate a striking chromosome dependence in scans with the full-length t1519 probe. The abundance of t1519 sites was found to be high in five chromosomes (16-18, 20 and Y), but low in all the others. Counts varied from 500 (chromosome 16) to 1429 (chromosome 18) in the high abundance group and 0 (7 chromosomes) to 22 (X chromosome) in the low (Table 1). Sub-region probes yielded matches in four chromosomes (15, 18, 19, and 21) with counts varying between 167 (sub-region 1 in chromosome 21) and 924 (sub-region 3 in chromosome 15). All chromosomes had at least one sequence match with one or more of the probes.

**Table 1:** Chromosome distribution of t1519 in protein-coding genes.

| chr | genes with<br>t1519 in high<br>abundance<br>chr | genes<br>with<br>t1519 in<br>other chr | genes with t1519 region<sup>a</sup> |  |  |  |  |
| --- | --- | --- | --- | --- | --- | --- | --- |
|  |  |  | 1 | 2 | 3 | 4 | 8 |
| 1 |  | 18 |  |  |  |  |  |
| 2 |  | 18 |  |  |  |  |  |
| 3 |  | 12 |  |  |  |  |  |
| 4 |  | 20 |  |  |  |  |  |
| 5 |  | 11 |  |  |  |  |  |
| 6 |  | 9 |  |  |  |  |  |
| 7 |  | 11 |  |  |  |  |  |
| 8 |  | 11 |  |  |  |  |  |
| 9 |  | 9 |  |  |  |  |  |
| 10 |  | 7 |  |  |  |  |  |
| 11 |  | 6 |  |  |  |  |  |
| 12 |  | 3 |  |  |  |  |  |
| 13 |  | 4 |  |  |  |  |  |
| 14 |  | 7 |  |  |  |  |  |
| 15 |  | 4 | 391 |  | 924 |  | 481 |
| 16 | 500 |  |  |  |  |  |  |
| 17 | 511 |  |  |  |  |  |  |
| 18 | 1429 |  |  |  |  | 568 |  |
| 19 |  |  |  |  | 344 |  |  |
| 20 | 589 |  |  |  |  |  |  |
| 21 |  |  | 167 | 260 |  |  |  |
| 22 |  | 1 |  |  |  |  |  |
| 23(X) |  | 22 |  |  |  |  |  |
| 24(Y) | 655 |  |  |  |  |  |  |

<sup>a</sup>: 180bp regions of t1519 sequence beginning at the 5' end.

### Location of t1519 sequence matches in the high abundance human chromosomes

The locations of t1519 sequence matches in the five high abundance chromosomes were downloaded from NCBI BLAST and plotted with Integrated Genome Viewer. The results showed that t1519 matches are found throughout the length of each chromosome (Fig. 3). Exceptions occur in the case of the Y chromosome where matches were found only in the leftmost 18Mb of the sequence, and in chromosome 16 where a large gap was found between positions ∼30Mb and 50Mb. t1519 matches were absent from the centromere in each of the five chromosomes.

**Fig. 3:**
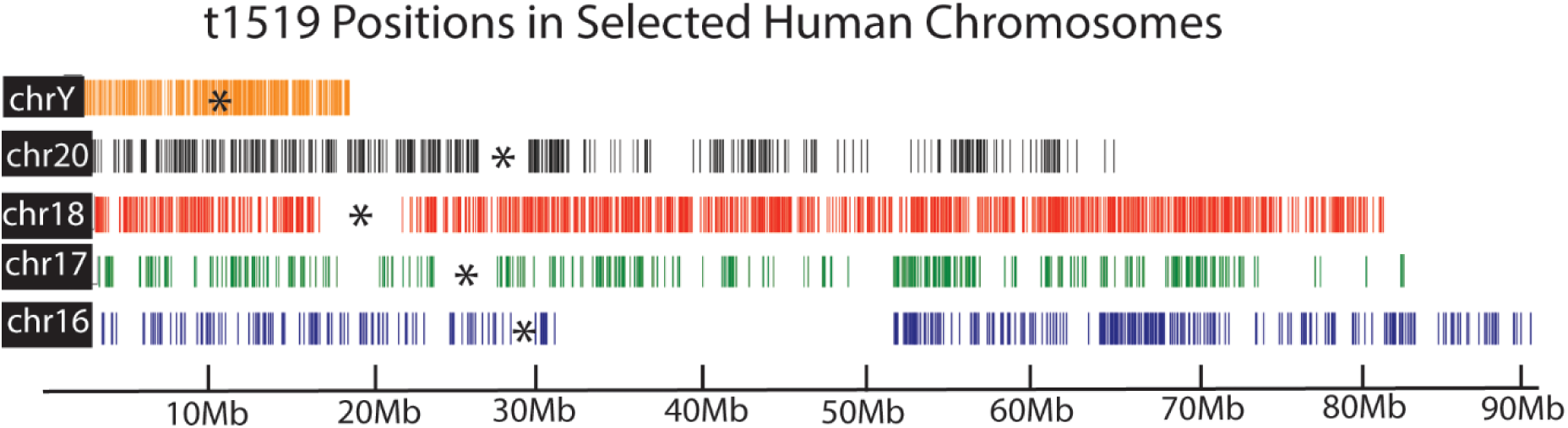
Plot of t1519 locations in the sequences of the five high abundance human chromosomes. Note that with only a few exceptions, t1519 matches are distributed along the entire length of the five chromosomes.

### Identification and comparison of t1519-containing genes with control genes lacking t1519

Individual t1519-containing and control genes for analysis were identified by visual inspection of sequence maps for three of the five high abundance chromosomes, chromosomes 16, 17 and 18 (Fig. 3). Matches were all found within introns. Expression level was plotted against length for each gene. Separate analyses were carried out for each chromosome.

### Protein-coding genes

Results for chromosome 16 genes with and without t1519 are shown in Figs. 4a and 4b, respectively. Comparison of the two shows a clear correlation of t1519 with long genes. Nearly all genes greater than ∼200Mb in length contain t1519, while genes of the same length but lacking t1519 were not observed. In contrast, genes in the common length range (less than ∼200Mb) were observed either with or without t1519. A quantitative correlation involving t1519 elements was observed with genes in the common length range. Average expression levels were 47.1 ± 51.1 TPM (n=76) and 90.1 ± 133.5 TPM (n=80), respectively in t1519 containing and those lacking t1519. The result suggests t1519 may have the expected suppressive effect on expression of common length genes. A quantitative correlation involving long gene expression was also observed. Expression of long genes was found to be limited to ∼50 TPM while much higher levels (up to ∼600 TPM) were the rule among genes with lengths in the common range.

**Fig. 4:**
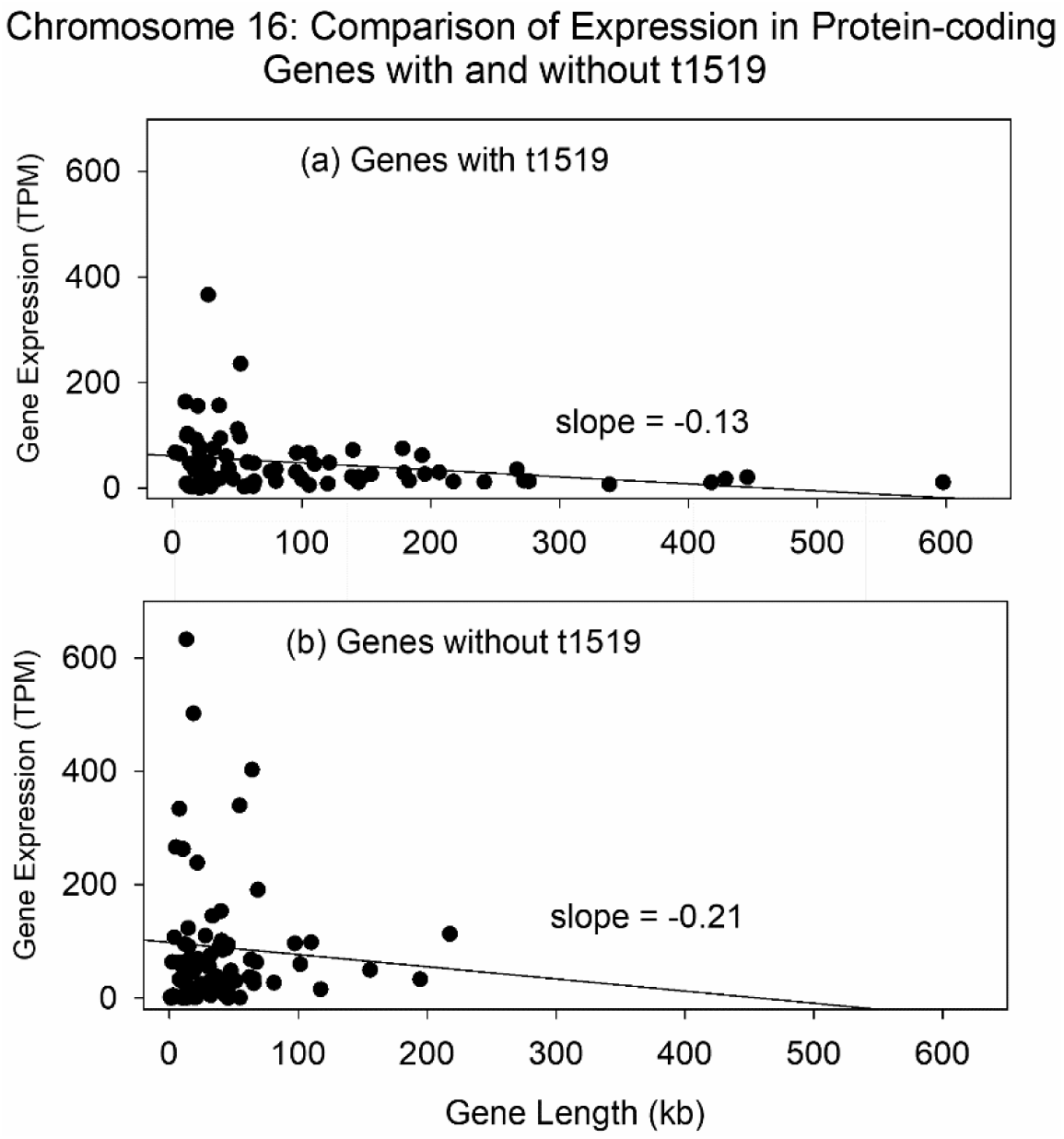
Plot of gene expression against gene length for chromosome 16 protein coding genes with (a) and without (b) t1519 elements within the coding region. Note the absence of long genes lacking t1519. Data for this plot can be found in Supplementary Table S5.

Findings with chromosomes 17 and 18 (Figs 5 and 6) were similar to those described above for chromosome 16. In chromosomes 17 and 18 long gene expression was found only in t1519-containing genes. Long genes lacking t1519 were not observed. Genes in the common length range could have t1519 or not, but those with t1519 had a lower average expression level. In chromosome 17 average expression levels were 35.6 ± 43.1 TPM (n=58) and 56.8 ± 48.7 TPM (n=55), respectively for t1519-containing and control genes. For chromosome 18 genes the comparable values were 40.4 ± 66.6 TPM (n=102) and 48.0 ± 72.3 TPM (n=95). As in the case of chromosome 16 long genes, long gene expression in chromosomes 17 and 18 was limited to a low expression level of ∼50 TPM.

**Fig. 5:**
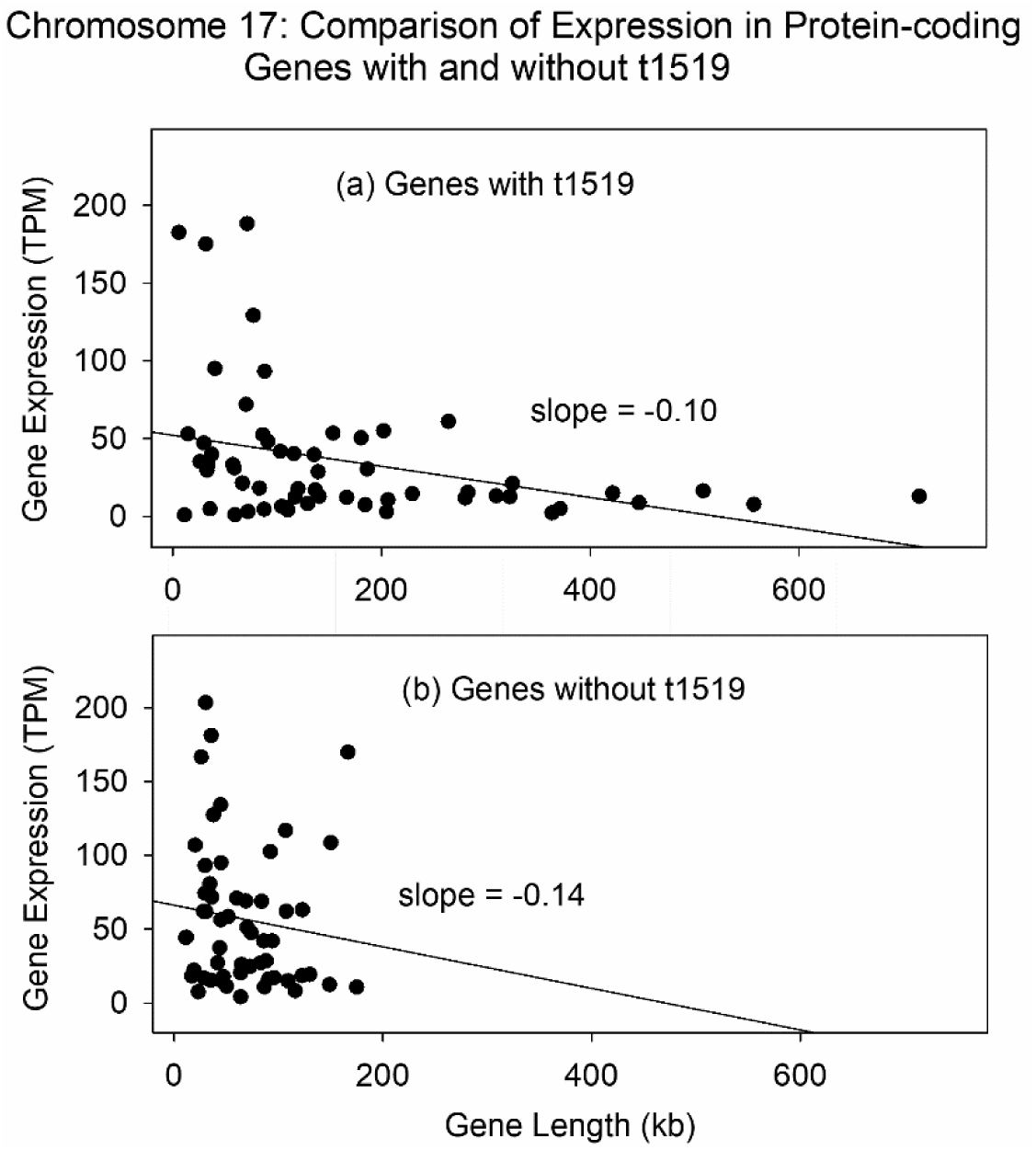
Plot of gene expression against gene length for chromosome 17 protein coding genes with (a) and without (b) t1519 elements within the coding region. Note the absence of long genes lacking t1519. Data for this plot can be found in Supplementary Table S6

**Fig. 6:**
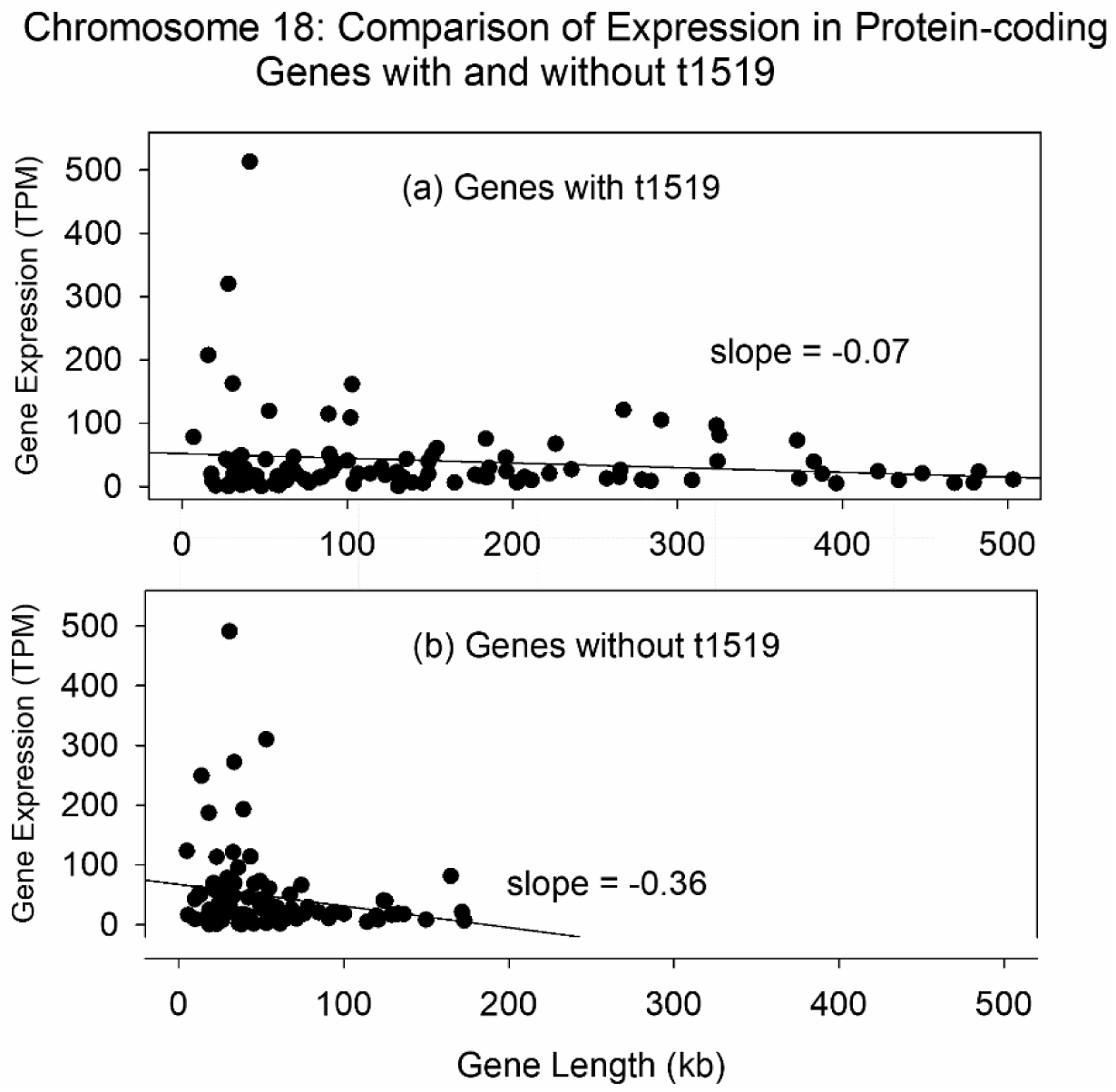
Plot of gene expression against gene length for chromosome 18 protein coding genes with (a) and without (b) t1519 elements within the coding region. Note the absence of long genes lacking t1519. Data for this plot can be found in Supplementary Table S7.

The observed correlation of t1519 with long genes was not expected. The result suggests t1519 provides a function that enables long gene expression but is not required for expression of genes in the common length range. One possibility is that t1519 may have a strong effect to counteract transcriptional delays, an effect also demonstrated for the mouse gene Sfpq [17–18].

The observed suppressive effect on expression of genes in the common length range was expected. The idea that “extra” transcriptional termination signals in L1 could antagonize gene expression by causing premature transcription termination can account for the observations reported here [4–6]. Finally, the effect of t1519 to constrain long gene expression to a low range is a novel finding. The observation may be due to a combination of distinct activating and suppressive effects of t1519 on long gene expression [4, 19–21].

### lncRNA genes

Comparison of lncRNA genes with and without t1519 was carried out using the same strategy described above for protein-coding genes. Populations of t1519-positive and negative genes were first identified in each of the three high-t1519 abundance chromosomes. Expression level was then plotted against length for each gene and the results were compared between t1519-positive and negative populations (Figs 7-9).

**Fig. 7:**
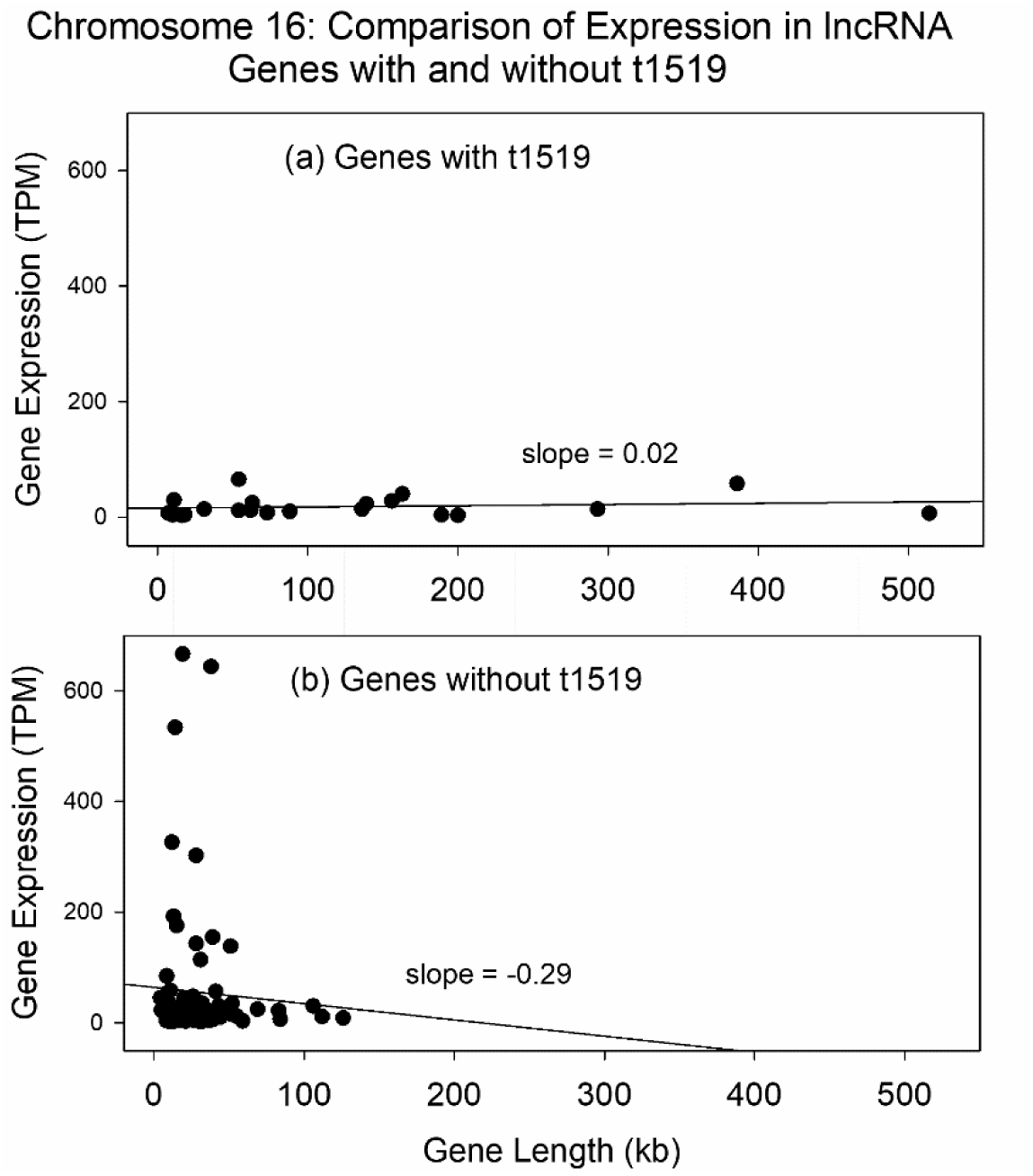
Plot of gene expression against gene length for chromosome 16 lncRNA genes with (a) and without (b) t1519 elements within the coding region. Note the absence of long genes lacking t1519. Note also the weak expression of both long and common length genes containing t1519 elements. Data for this plot can be found in Supplementary Table S8

The results showed the same correlation of t1519 with long lncRNA genes observed with long protein coding ones. All long lncRNA genes contained at least one t1519 sequence while no long lncRNA genes lacking t1519 were found. Among genes in the common length range, t1519 was found to correlate with suppression of expression. The effect was particularly strong in chromosomes 16 and 17 (Figs 7 and 8) although a modest effect was also observed with chromosome 18. Average expression values in t1519-positive and negative genes were: chromosome 16: 18.2 ± 17.6 TPM (n=21), 55.9 ± 123.5 TPM (n=84); chromosome 17: 138.7 ± 227.3 TPM (n=28); 292.0 ± 482.4 TPM (n=98); chromosome 18: 55.6 ±105.0 TPM (n=79); 63.0 ± 102.4 TPM (n=29). As observed with protein-coding genes, the expression level of lncRNA genes varied only modestly over the length range that long genes differ from common length ones (Figs 7-9). A low level of expression was observed in genes throughout the long gene range. This observation indicates that while t1519 correlates with an effect to enable long gene expression, the effect supports only a low level of expression.

**Fig. 8:**
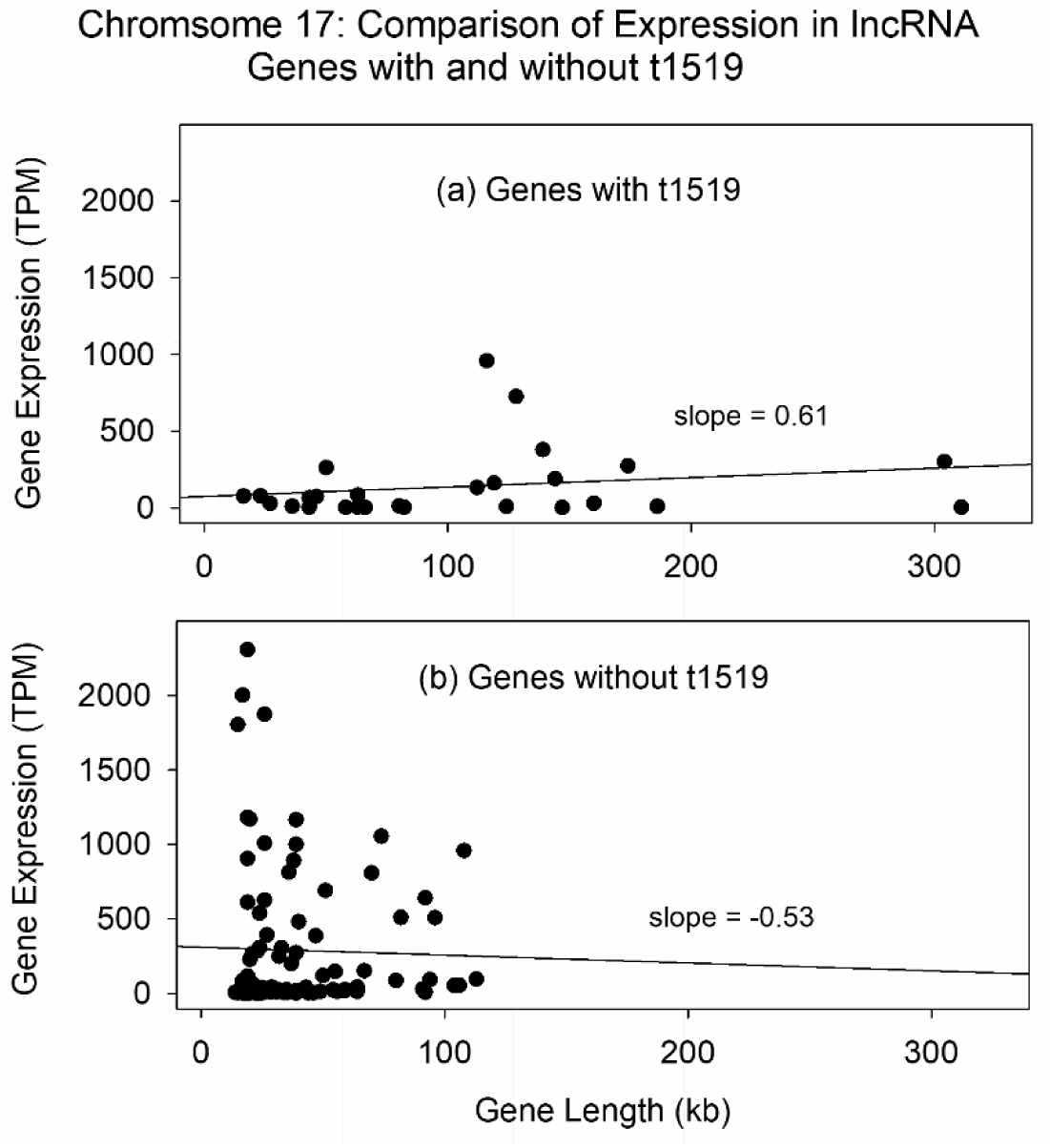
Plot of gene expression against gene length for chromosome 17 lncRNA genes with (a) and without (b) t1519 elements within the coding region. Note the absence of long genes lacking t1519. Note also the weak expression of both long and common length genes containing t1519 elements. Data for this plot can be found in Supplementary Table S9.

**Fig. 9:**
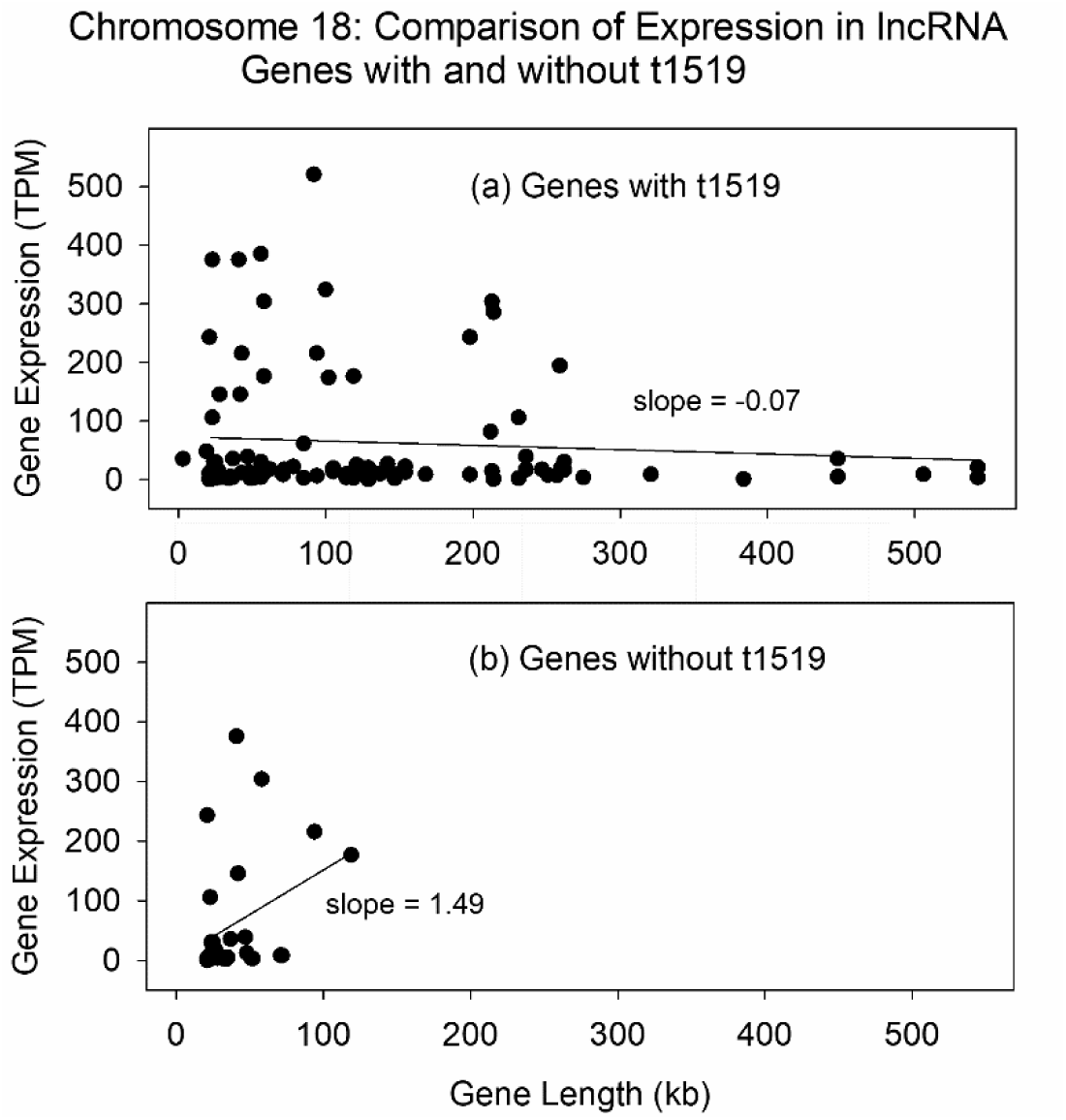
Plot of gene expression against gene length for chromosome 18 lncRNA genes with (a) and without (b) t1519 elements within the coding region. Note the absence of long genes lacking t1519. Note also the strong expression of both long and common length genes containing t1519 elements. Data for this plot can be found in Supplementary Table S10.

### Multiple t1519 sequences in the same gene

An important feature of the results reported here has to do with the fact that many of the genes examined have more than one t1519 sequence within the coding region. Genes with more than one t1519 are found in both protein-coding and lncRNA genes and also among both long and common length genes. Genes with multiple t1519 elements were examined further by plotting t1519 count against gene length in protein-coding and lncRNA genes of chromosomes 16, 17 and 18.

The results with chromosome 16 protein-coding genes illustrate the basic findings (Fig. 10a). A positive correlation between t1519 count and gene length was observed throughout the range of gene length examined. A similar positive correlation was found with protein-coding genes in chromosomes 17 and 18 (Fig. 10 b and c) and with all lncRNA genes (Fig. 11 a-c). The results support the idea that any effect of t1519 on gene expression would be one that benefitted from repetition of t1519 elements as RNA synthesis proceeds along the DNA template. An alternative explanation is that each t1519 insertion event has been preserved during evolution because of its ability to downregulate expression of the host gene. According to this interpretation, the group of insertions would constitute a historical record of changes in the host gene’s level of expression.

**Fig. 10:**
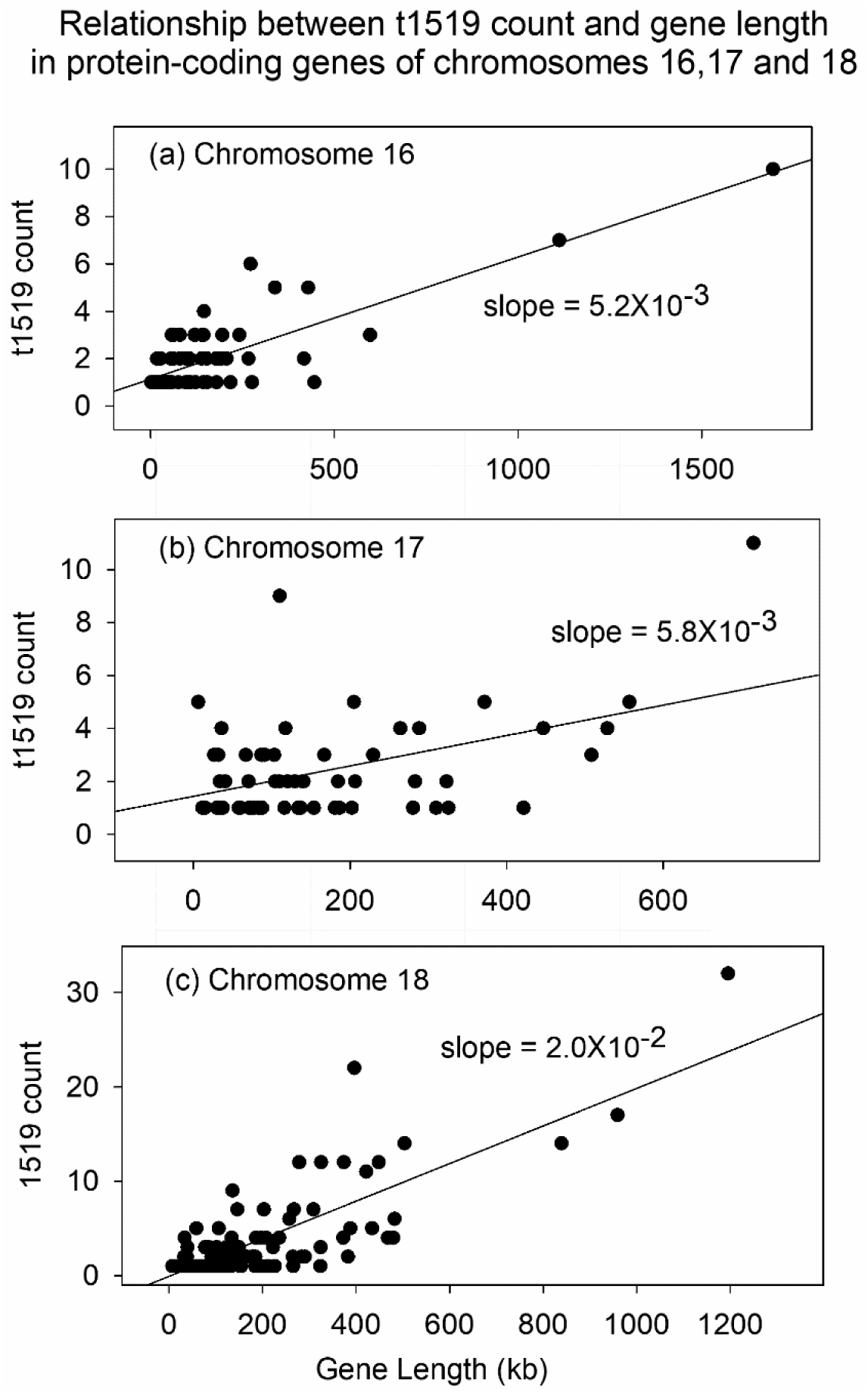
Plots of t1519 count against gene length for protein-coding genes of chromosomes 16, 17, and 18. Note that a positive slope is observed in all three plots.

**Fig. 11:**
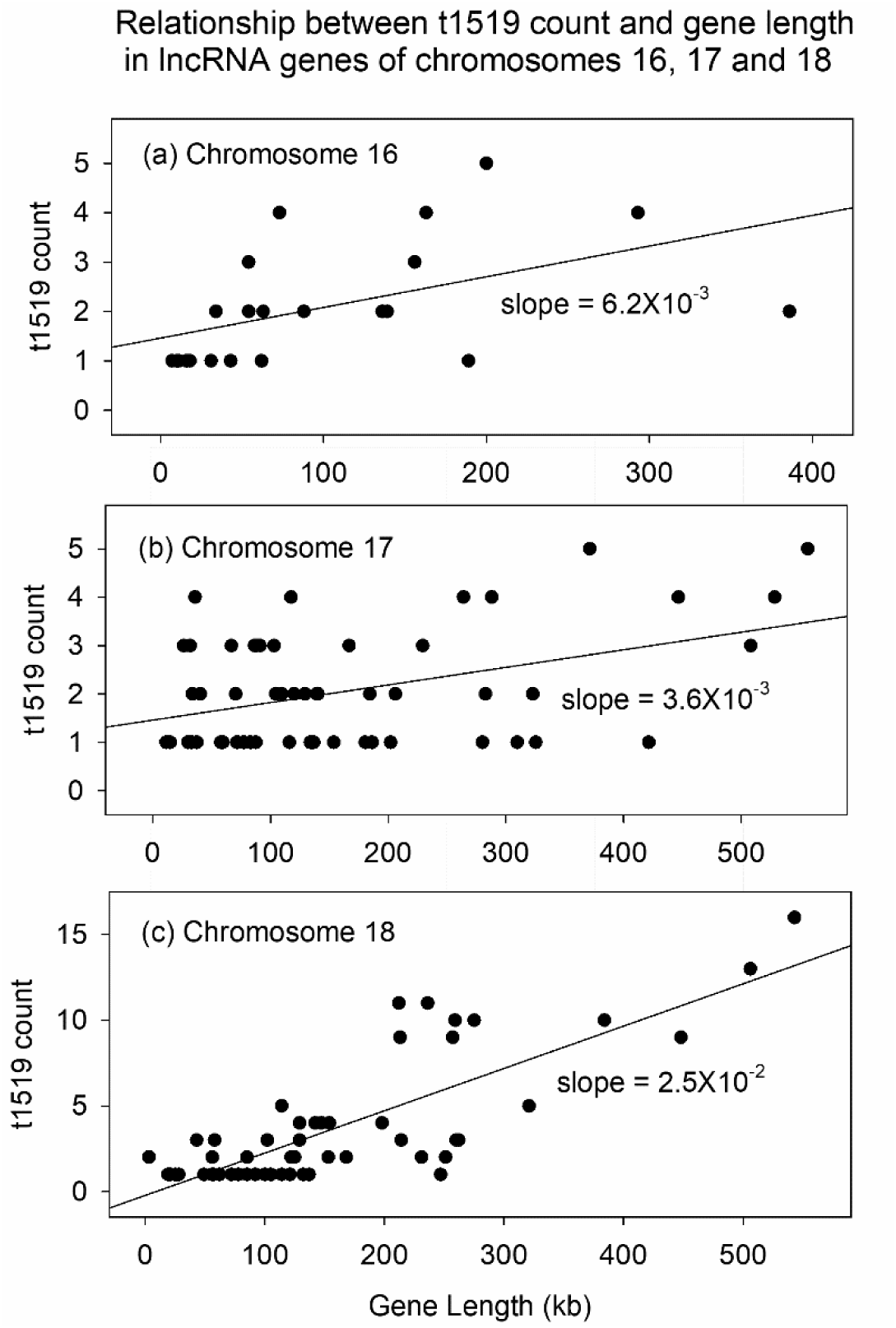
Plots of t1519 count against gene length for lncRNA genes of chromosomes 16, 17, and 18. Note that a positive slope is observed in all three plots

## Discussion

The results reported here relate to the relationship between gene length and gene expression in the human genome [22]. Two types of genes can be recognized: (1) those in the common range with lengths of up to ∼140kb; and (2) long genes with lengths greater than ∼140kb. The level of gene expression is distributed quite differently in the two populations. Most human genes (85%-90%) have lengths in the common range, and the level of expression is a negative function of gene length. Short genes have a high level of expression and long common length genes have a low level. In contrast, expression of most long genes is distributed over a narrow window at the low level of the overall expression range (<50 TPM). The two distribution types are illustrated in Figure 5, which shows a plot of gene expression against length for protein-coding genes in chromosome 17. Figure 5b illustrates the plot characteristic of genes in the common length range. All genes are less than ∼140kb in length and a negative slope is observed relating expression to gene length. The negative slope is thought to result, at least in part, from the fact that more short transcripts can be synthesized in the same time as fewer longer ones.

Figure 5a illustrates the distribution of expression levels found in long genes. In all genes are greater than ∼140kb in length the expression level is low (< 50 TPM). The expression level remains low over a broad range (400kb or more) of genome length. It is not known how the expression level of long genes is constrained to the narrow window observed.

### Long genes

The findings reported here relating to long genes show that t1519 is present in the body of all long genes in human chromosomes 16-18. The finding applies to both protein-coding and lncRNA genes. By enabling long gene expression, t1519 and other factors allow humans to benefit from the extra information encoded in long compared to common length genes. It is not known how t1519 may act to enable long gene expression, but information is available about other factors that have the same effect. Mouse Sfpq, for instance, is found to potentiate long gene expression by maintaining the transcription complex in an active state [17–18]. Spt5 functions to favor long gene expression by preventing detachment of the RNA polymerase II complex from the DNA template as transcription proceeds [23]. Topoisomerases favor long gene expression by preventing template DNA overwinding [9, 24–25]. A variety of mechanisms have been demonstrated to antagonize transcriptional pauses, mechanisms that have a preferential effect on long gene expression [11].

In addition to the effect of t1519 to enable long gene expression, the presence of t1519 also correlates with the way long gene expression is confined to a low level. Examples of this effect can be seen, for instance, in Figures 5 and 8. It is suggested that confining long gene expression in this way would prevent expression of long genes from causing the cell to exceed its overall capacity for transcription activity. While the mechanism of transcription inhibition in long genes has not been reported, inhibition by premature termination would appear to be an attractive possibility [4–6].

### Genes in the common length range

t1519 was found in the body of genes in the common length range as well as in long genes. In contrast to the situation with long genes, however, t1519 is found to be present in some common length genes, but not others. This distribution is observed in both protein-coding and lncRNA common length genes. Further, in the expression vs. gene length plots reported here, the presence of t1519 is found to correlate with attenuation of gene expression in every population examined. Attenuation may be quantitatively strong as in the case of lncRNA genes in chromosomes 16 and 17 (Figs 7 and 8) or more modest as in chromosome 18 protein coding genes (Fig 6), but activation was not observed in any population. The observation of widespread suppression is consistent with the idea that t1519 insertions act primarily by their ability to cause premature termination of transcription as mentioned above.

### Distribution of t1519 among human chromosomes

No explanation is available for the puzzling distribution of t1519 elements among the human chromosomes. Hundreds of t1519’s are found in each of the five high abundance chromosomes (i.e. 16-18, 20 and Y), but many fewer are found in the others (see Table 1). Why these five chromosomes? How do long genes in non-high abundance chromosomes get along without the functions of t1519 documented here? Could another element or elements supply the same functions? One possible explanation involves the natural history of L1 elements. If the source or targets of retro-transposition events were to change over evolutionary time, then in the past another feature could have provided the same functions as t1519 provides today. Such events might have escaped the analysis described here. Clarifying the nature of such alternative elements might be revealing about the nature and evolution of the genome.

## Acknowledgements

I gratefully acknowledge the assistance of Anahita Golchin in the early stages of this investigation.

## Supplementary Material

Supplementary Table S1: DNA sequence of the t1519 element. https://github.com/micro456/line1/blob/main/t1519_sequence1.txt

Supplementary Table 2: Location of t1519 elements in human chromosome 16. https://github.com/micro456/line1/blob/main/chr16_t1519.bed.txt

Supplementary Table S3: Location of t1519 elements in human chromosome 17. https://github.com/micro456/line1/blob/main/chr17_t1519.bed.txt

Supplementary Table S4: Location of t1519 elements in human chromosome 18. https://github.com/micro456/line1/blob/main/chr18_t1519.bed.txt

Supplementary Table S5: Gene name, expression level, length and t1519 count for plots shown in Figures 4 and 10. https://github.com/micro456/line1/blob/main/chr16_all_prot_coding_genes2.xlsx

Supplementary Table S6: Gene name, expression level, length and t1519 count for plots shown in Figures 5 and 10. https://github.com/micro456/line1/blob/main/chr17_all_prot_coding_genes1.xlsx

Supplementary Table S7: Gene name, expression level, length and t1519 count for plots shown in Figures 6 and 10. https://github.com/micro456/line1/blob/main/chr18_all_prot_coding_genes1.xlsx

Supplementary Table S8: Gene name, expression level, length and t1519 count for plots shown in Figure 7 and 11. https://github.com/micro456/line1/blob/main/chr16_lncRNA_yes_no1.xlsx

Supplementary Table S9: Gene name, expression level, length and t1519 count for plots shown in Figures 8 and 11. https://github.com/micro456/line1/blob/main/chr17_lncRNA_yes_no1.xlsx

Supplementary Table S10: Gene name, expression level, length and t1519 count for plots shown in Figures 9 and 11. https://github.com/micro456/line1/blob/main/chr18_lncRNA_yes_no1.xlsx

